# Systematic biases in disease forecasting - the role of behavior change

**DOI:** 10.1101/349506

**Authors:** Ceyhun Eksin, Keith Paarporn, Joshua S. Weitz

## Abstract

In a simple susceptible-infected-recovered (SIR) model, the initial speed at which infected cases increase is indicative of the long-term trajectory of the outbreak. Yet during real-world outbreaks, individuals may modify their behavior and take preventative steps to reduce infection risk. As a consequence, the relationship between the initial rate of spread and the final case count may become tenuous. Here, we evaluate this hypothesis by comparing the dynamics arising from a simple SIR epidemic model with those from a modified SIR model in which individuals reduce contacts as a function of the current or cumulative number of cases. Dynamics with behavior change exhibit significantly reduced final case counts even though the initial speed of disease spread is nearly identical for both of the models. We show that this difference in final size projections depends critically in the behavior change of individuals. These results also provide a rationale for integrating behavior change into iterative forecast models. Hence, we propose to use a Kalman filter to update models with and without behavior change as part of iterative forecasts. When the ground truth outbreak includes behavior change, sequential predictions using a simple SIR model perform poorly despite repeated observations while predictions using the modified SIR model are able to correct for initial forecast errors. These findings highlight the value of incorporating behavior change into baseline epidemic and dynamic forecast models.

---

Infectious disease outbreaks catalyze public health responses. The scope of responses are increasingly tied to disease “forecasts”. Disease forecasts include estimates of infected cases, and associated mortality and morbidity^1^. Accurate forecasts can help specify when, where, and how much response is needed to control an outbreak, e.g., whether for SARS, Ebola or Zika. However, inaccurate forecasts can lead to a mismatch between resources needed vs. those deployed. Inaccurate forecasts can also diminish public support for future interventions^2,3^.

Dynamic models underlie the science of disease forecasting^4,5^. The forecasting methodology involves fitting a dynamic model to available data and then extrapolating forward in time (and sometimes in space and in time)^6–8^. The extrapolation step depends on assumptions concerning subsequent disease transmission and recovery. These assumptions are informed, in part, by the initial fitting. Individual models may differ with respect to quantitative details, e.g., the probability of infection given contact, and even qualitative details, e.g., whether certain modes of transmission are possible. Such differences can underlie variation in predictions, particularly for emerging outbreaks in which prior information on transmission may be limited^9,10^.

Despite their differences, leading model-based forecasting efforts often base their predictions on the assumption that individuals continue to behave in the same way irrespective of disease prevalence^10^. Given this assumption, it follows that the initial “speed” at which cases increase is informative with respect to the cumulative “size” of the outbreak. For example, if a disease spreads faster initially in outbreak A than in outbreak B, then more people will be infected in A who themselves continue to spread the disease at a faster rate than in B, all else being equal in the two outbreaks. In the long-term this should lead to a strong correlation between the initial speed and the final size (high in A, low in B).

However, the speed-size relationship may become tenuous when accounting for changes in individual behavior during an outbreak^11^. Changes can include social distancing and other protective measures. For example, during the 2009 H1N1 pandemic, 59% to 67% of Americans washed their hands more frequently, more than 50% were prepared to stay at home if they or a member of their family were to get sick, and 25% avoided crowded areas^12^. As another example, decisions to forego traditional burial ceremonies during the Ebola virus disease (EVD) outbreak in West Africa reduced the rate of post-death transmission of EVD^13–15^. Such behavior change can significantly reduce disease transmission even in the absence of pharmaceutical interventions ^16–19^. Yet, the consequences of changing behavior are generally not accounted for in the science of disease forecasting. In this paper, we provide a model-based rationale for monitoring behavior change during the initial stages of an outbreak to avoid predictable pitfalls of forecasts.

## Results

### The influence of behavior change on final outbreak size

We modify the standard SIR model to include a social distancing term *a* that is a function of the number of infected *I* and recovered *R*. See Methods section for model details. The social distancing term scales the transition rate from a susceptible state to an infected state, that is given by *aβSI*/*N* where *β* is the infection rate, *S* is the number of susceptible individuals, *I* is the number of infectious individuals and *N* is the population size. We consider two types of social distancing models. In the first model, individuals reduce their interaction with others proportional to the cumulative fraction of affected (infectious and recovered (*I* + *R*)/*N*) individuals, i.e., *a* = (1 − (*I* + *R*)*/N*)^*k*^. We term this “long-term awareness”. In the second model, the reduction is proportional to the percentage of infectious individuals (*I*) at a given moment, i.e., *a* = (1 − *I*/*N*)^*k*^. We term this “short-term awareness”. When the average infection in the population is high, the social distancing action is close to zero implying individuals are taking the utmost preemptive measures. In both distancing models, we raise the reduction term to the *k*th power for some behavior parameter *k* ≥ 0. When *k* = 0, we recover the benchmark SIR model with *no distancing* response to disease prevalence. The distancing model is *linear* when *k* = 1. For *k* > 1, the social distancing model is *nonlinear*. As the nonlinearity (*k*) increases, individuals become more sensitive to disease prevalence^20,21^.

The endemic level of the disease can be identified by finding the equilibrium of the model. We consider the long-term awareness model where distancing term is proportional to the fraction of affected. Assuming *R*(0) = 0 and *S*(0) ≈ *N*, and noting that *R*(∞) = *N* − *S*(∞) since *I*(∞) = 0, we obtain the following infinite time relationship for susceptible individual population, 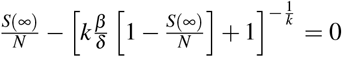 for *k* > 0 (see SI section for the derivation). Solving this equation when distancing is linear (*k* = 1), we have the fraction of recovered at equilibrium as 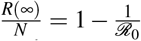 where *ℛ*_0_:= *β* / *δ* is the basic reproduction number. It is not possible to obtain a generic closed form solution for arbitrary *k*. In Figure 1, we plot the equation for *k* = {1,2,3}, and compare the solution to this equation for *S*(∞) with the infinite time relationship of the SIR model with no distancing (*k* = 0)^22^. As is evident, social distancing has the potential to significantly increase the fraction of uninfected individuals at the end of the epidemic relative to the SIR model without behavior change. We cannot provide a similar derivation of equilibrium points for the short-term awareness model, and turn instead to numerical simulations to measure asymptotic case counts for different models of short-term awareness.

**Figure 1.**
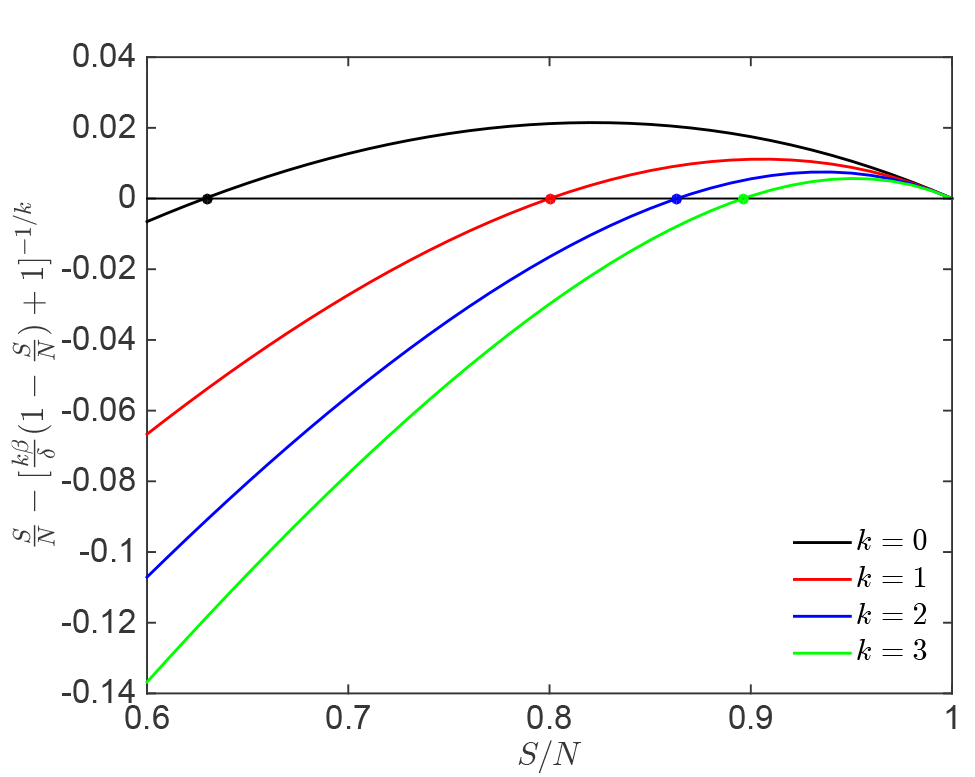
Proportion of susceptible individuals at equilibrium given SIR dynamics with behavior change. We let *ℛ*_0_ = *β* / *δ* = 1.25. For *k* = 0, we use the well known infinite time relationship for *S*(∞) of the standard SIR dynamics^22^ (see SI Section A). The non-trivial zero intersection of the SIR model with no distancing is at *S*(∞)/*N* = 0.63 and it is the equilibrium point. For *k* = {1,2,3} we plot the left hand side of infinite time relationship for *S*(∞)/*N*. The non-trivial zero intersection of the function is at 0.8, 0.86, and 0.9 when *k* is equal to 1, 2, and 3, respectively.

### The influence of behavior change on speed-size relationships

We define the speed of the disease outbreak in an SIR model with social distancing as the initial increase in infectious cases given a single infected individual in an otherwise susceptible population. Given the baseline dynamical systems, this speed is the infection rate (*β*) minus the healing rate (*δ*), *r* = *β* − *δ*, for both social distancing models because initially the number of susceptible individuals is approximately equal to the population size (*S*(0) ≈ *N*, *I*(0) ≈ 0 and *R*(0) = 0). The speed *r* has units of inverse time and can be rewritten as *r* = *δ* (*ℛ*_0_ − 1).

To compare the effects of short-and long-term awareness of speed-size relationships, we simulate the SIR dynamics with social distancing, given *ℛ*_0_ = *β* / *δ* = 1.25. We have *k* = 0 for the benchmark model without social distancing. For both short-and long-term awareness, we let *k* = 1 and *k* = 3 for the linear and nonlinear social distancing models, respectively. We plot the percentage affected ((*I*(*t*) + *R*(*t*))/*N*) from the disease with respect to time in Fig. 2. Note that since the eventual average number of infected is zero (*I*(∞) = 0), the eventual percentage affected is equal to *R*(∞)/*N* = 1 − *S*(∞)/*N*.

**Figure 2.**
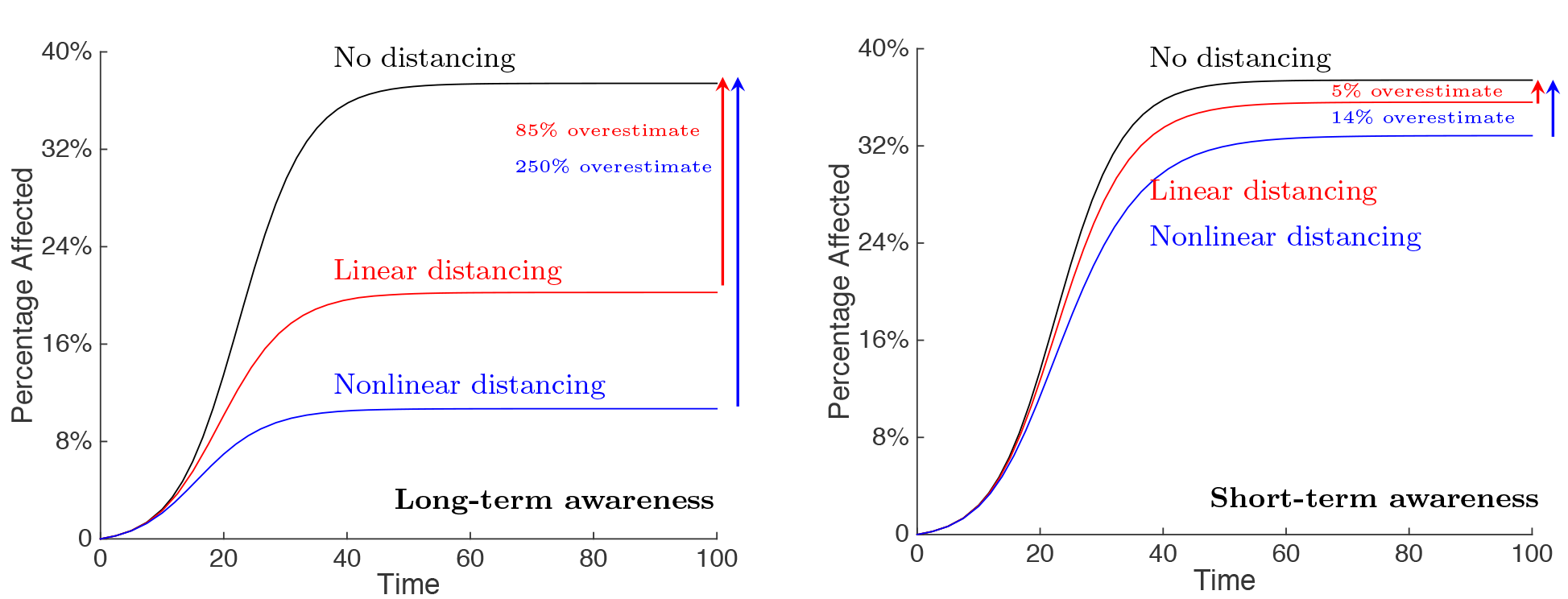
Effect of social distancing on disease severity forecasts. We show the spread of disease for each model with respect to time given a basic reproduction number of *ℛ*_0_ = 1.25. We consider two social distancing models including long-term and short-term awareness in the left and right panels, respectively. (left) The percentage affected when individuals distance with respect to total percentage of infected and recovered. (right) The percentage affected when individuals distance with respect to percentage of infected. At the beginning all models have comparable outbreak speeds. The simulation time horizon is 100. The benchmark model is the same in both figures given model parameters. See the Methods for details regarding the model.

Figure 2 demonstrates the potential pitfall of disease forecasting without accounting for changes in social behavior during an outbreak. The epidemic dynamics given distancing with long-term awareness is shown in Figure 2 (left). The size of the epidemic is significantly lower in the social distancing model than in the benchmark model. Counterintuitively, there is no early signal that would inform an *a priori* forecast of such a reduction. The initial *speed* at which cases increase at the outset of the outbreak is nearly the same in both scenarios. That is to say, the number of cases increases as fast in the social distancing model as in the benchmark model. A forecast that leverages the initial speed to gain traction on disease parameters but assumes no behavior change would overestimate the final size by 85%.

The basis for this discrepancy in Figure 2(left) can be understood by recognizing that the fraction of infectious individuals is small at the early stages of an outbreak. In a linear social distancing model, such a fraction is too small to substantively change individual behavior. As the outbreak grows in size, the fraction of infectious individuals is large enough to influence individual behavior. When individuals decrease their infectious contacts, either via reducing interactions or by taking protective measures during risky interactions, the disease spreads slower than the benchmark model. Eventually, far fewer individuals are infected than would have been predicted via a standard extrapolation approach. In Figure 2(left) we also show that the over-estimation of spread can be much higher (250% in the example) when individual response to disease prevalence is nonlinear.

When individuals respond to the prevalence of the infected, they respond to the current threat only. Figure 2 (right) shows the resulting epidemic dynamics given distancing with short-term awareness. As is evident, the error in over-estimation is smaller (5%) when individuals distance with respect to the present threat only rather than to the present and past threat. Furthermore, over-estimation of epidemic size increases modestly (to 14%) when the social distancing response is nonlinear given short-term awareness.

### Sequential forecasting of a SIR outbreak with behavior modification

The prior results suggest that early forecasts that do not account for behavior change may overestimate final epidemic size. Further, these forecasts are prone to further errors given uncertainty in disease spread parameters and initial state of the disease. In practice, early forecasts are updated during an outbreak, given measurements of incidence and iterative model updating^7^. In this section, we use an ensemble-based Kalman filter (EnKF), described in the Methods, to analyze the expected error in sequential forecasts that leverage new information on disease spread given a baseline outbreak with behavior change.

We consider two sequential forecasting scenarios based on noisy measurements of the fraction of infected. In the first scenario, EnKF predictions do not account for behavior change making predictions based on SIR dynamics with the underlying behavior response *a* = 1. That is, the behavior parameter is assumed to be zero 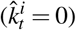 and the state estimate by the *i*th ensemble member is given by 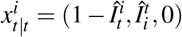 for all *t* where we recall *x*_*t*_ = (*S*_*t*_/*N*,*I*_*t*_/*N*,*k*). In the second scenario, EnKF predictions treat behavior parameter *k* as a random variable. Specifically, state realizations of *k* come from a Gaussian prior distribution with mean 1 and variance equal to 0.5, i.e., we let 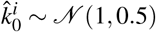. The *i*th state realization is given by 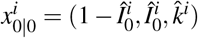.Then the behavior parameter estimate is updated after each observation.

We first generate an epidemic outbreak given long-term awareness (Figure 3(a)-(b)). Then, we use the EnKF method to predict trajectories given weekly updates assuming an underlying model without (first scenario) and with behavior (second scenario) modification (Figures 3 (a) and 3 (b), respectively). Both models start with inaccurate predicted trajectories where the initial mean trajectories overestimate the ground truth trajectory. Further, the predictions using the model with variability in *k* has a higher initial cone of uncertainty. The predictions using the model with variability quickly increase in accuracy, i.e., the mean predictions become close to ground truth and cone of uncertainty shrinks, whereas the model without behavior continues to have inaccurate mean predicted trajectories, and does not give indication that they are inaccurate, i.e., the cone of uncertainty disappears.

**Figure 3.**
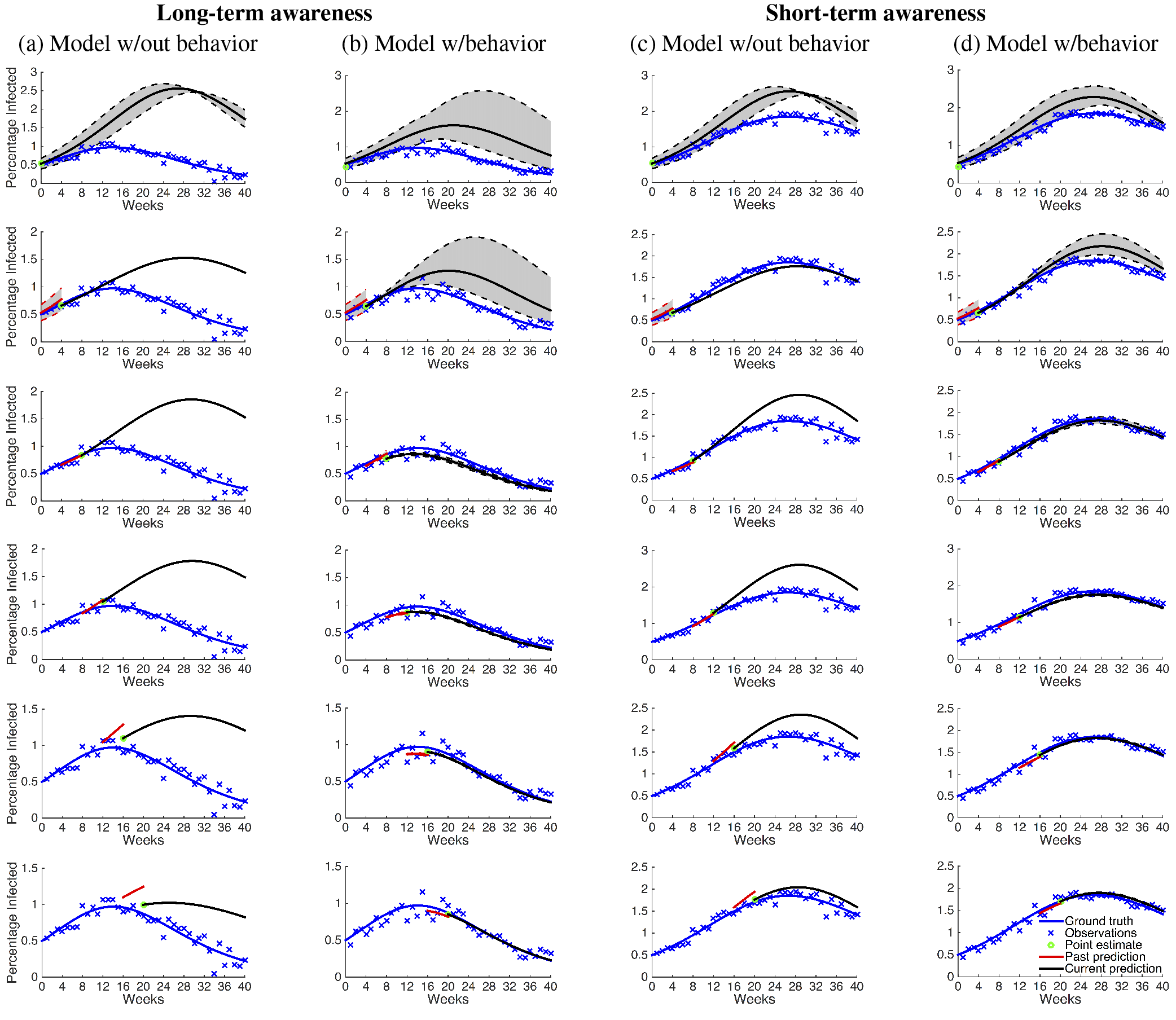
Sequential prediction of a synthetic SIR outbreak with EnKF. The blue lines show the course of the synthetic outbreak. Ground truth outbreak is generated using SIR dynamics with nonlinear distancing (*k* = 3). In columns (a)-(b), distancing model is long-term awareness. In columns (c)-(d), distancing model is short-term awareness. We mark the measurements of fraction of infected with ‘x’. The green circles show the posterior mean estimates of the fraction of infected *x*̄_*t*|*t*_ for the month in consideration. (a) and (c) show monthly predictions based on an inaccurate SIR model (*k* = 0), i.e., assuming *a* = 1. (b) and (d) show monthly predictions based on an accurate SIR model that accounts for behavior change. For (a) and (c), prediction noise distribution is zero mean Gaussian with variance 10^−10^. For (b) and (d), prediction noise distribution is zero mean Gaussian with variance 10^−4^—see SI Section C for details on how to select the variance. In all figures, black lines show the predicted trajectories of the EnKF starting from the current week. Red lines show mean predictions of the previous month. Shaded areas show the range of the trajectories in the ensemble. EnKF predictions that do not account for behavior change ((a) and (c)) perform poorly in comparison to the EnKF predictions that do ((b) and (d)).

Next, we generate an epidemic outbreak given short-term awareness (Figure 3(c)-(d)). Then, we use the EnKF method to predict trajectories given weekly updates for the first and second scenarios (Figures 3 (c) and 3 (d), respectively). The predicted trajectories follow a similar trend as in the long-term case. That is, the model with behavior changes provide accurate forecasts starting from the second month while the model without behavior consistently overestimate the ground truth trajectory. Here, forecast errors for the model without behavior changes are smaller because the short-term awareness model has a weaker influence on the ground truth trajectory than the long-term awareness model.

### Sequential learning of behavioral response during an outbreak

The results above show that in both long-term and short-term awareness models, sequential forecasts with an underlying model that consider behavior change are able to predict the future trajectory of the disease after enough observations. Here, we investigate whether the accurate predictions are a consequence of learning the true value of the behavior parameter. Figs. 4 (left) and (right) show the time evolution of the estimate of the behavior parameter 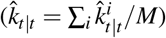for the long-term (Fig. 3(b)) and short-term awareness (Fig. 3(d)) distancing scenarios, respectively. We observe that both the mean estimates 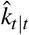 and the range of ensemble values for *k* approach the true behavior parameter value (*k* = 3). This shows that the sequential forecasts with EnKF are able to learn the behavior parameter, and hence accurately forecast the future trajectory.

**Figure 4.**
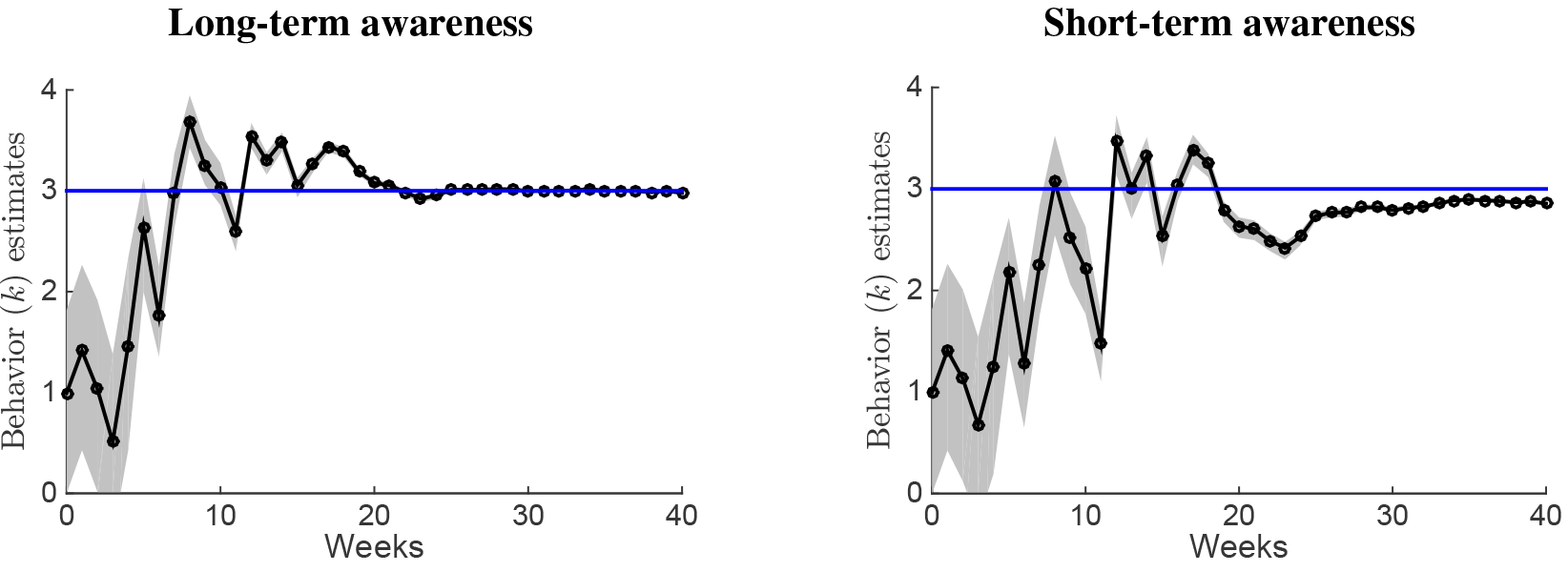
Sequential behavior parameter (*k*) estimates. Blue line shows the ground truth value of the behavior parameter *k* = 3. (Left) shows the improvement of the mean estimate of the behavior parameter *k̅* for the second scenario in the long-term awareness model (Fig. 3(b)). (Right) shows the improvement of the mean estimate of the behavior parameter *k̅* for the second scenario in the short-term awareness model (Fig. 3(d)). The shaded regions show the range of the *k* values in the ensemble. The ensemble values converge toward each other as more measurements are collected.

## Discussion

Our results point to a potential pitfall in the science of disease forecasting. Despite successes for respiratory diseases^23–26^, the accuracy of disease forecasts has been questioned-particularly in light of the gap between the realized and predicted number of cases for the EVD outbreak in West Africa in 2014-2015^2^. On the one hand, disease forecasts can provide a road map for prevention of a bad outcome through a targeted intervention^3^. Nonetheless, the road map should include an accurate baseline of what might occur if no interventions were implemented^19^. Ensuring that resources deployed include a buffer to achieve desired outcomes is essential. Prevention and control campaigns must also weigh the costs of over-allocation for resource management and logistics. The substantive differences in outcomes given social distancing based on long-vs. short-term awareness also present an opportunity to improve predictions. We suggest there is a critical need to systematically study and collect behavioral information on the “stickiness” of factors-including present and past infections-that inform individual decisions to modify behavior during an outbreak.

Our results point to potential opportunities in accounting for behavior change during sequential forecasts. In particular, data assimilation methods can help us learn about behavior change during an outbreak based on seemingly unrelated data such as number of infected, or new incidences. Indeed, including behavior change in our forecasting models can prove valuable in making sense of new data. In contrast, neglecting behavior change can result in inaccurate forecasts. Despite the promise, learning behavior-related parameters requires accurate measurements, particularly during early stages of an outbreak (see Fig. 3, Fig. 4, and SI). In addition, the success of the EnKF method depends on selection of parameters which were pre-assigned given the synthetically generated outbreak data. How to select the prediction noise in the absence of ground truth is an open research question (see SI Section C for additional details).

Here, we have provided theoretical and numerical evidence in support of reconsidering the accuracy of disease forecasts at the outset of an outbreak. We argued that comparative predictions of final sizes should not necessarily be predicated on estimates of initial speeds. Instead, accurate extrapolations from speed to size depend critically on changes in behavior. While our focus here was on individual behavior response to disease prevalence, individual behavior responses can take different forms during an outbreak, e.g., replacement of infected essential personnel with healthy ones^27^, and they can depend on seemingly exogenous phenomena, e.g., weather or co-infections (flu and pneumonia)^28^. An interdisciplinary research effort is necessary to understand the coupling between intrinsic changes in behavior and public health efforts to modify individual behavior before, during, and after an outbreak.

## Methods

### SIR model with social distancing

We modify the standard SIR model to include a social distancing term *a*(*I, R*) which is a function of the number of infected *I* and recovered *R*,

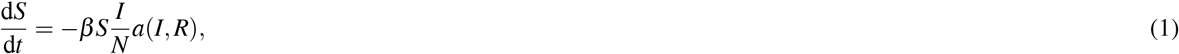

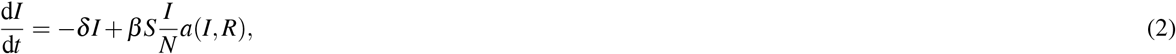

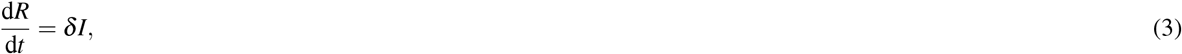

where *β* and *δ* are the disease infection and healing rates respectively, and *N* = *S* + *I* + *R* is the population size. The social distancing term (*a*(*I*, *R*): ℝ^2^ → [0, 1]) slows transition from a susceptible state to an infected state.

In the “long-term awareness” model, the social distancing term decreases proportional to the cumulative fraction of affected (infectious and recovered) individuals,

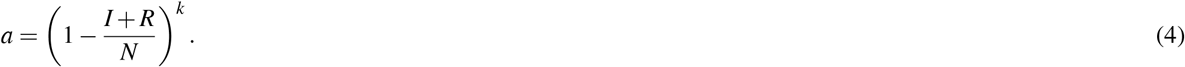

In the “short-term awareness” model, reduction is proportional to the percentage of infectious individuals

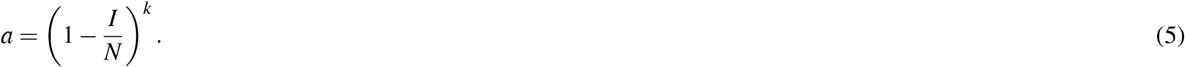

In numerical simulations, we use outbreaks generated using the SIR model (1)–(3) with long-term (4) or short-term (5) awareness as ground truth. We will fix infection and healing rates to *β* = 0.5 and *δ* = 0.4 respectively. Throughout the paper, the initial state of the synthetic outbreak will be *S*_0_/*N* = 99.5%, *I*_0_/*N* = 0.5%, and *R*_0_/*N* = 0%. We consider both linear and nonlinear distancing models. In the nonlinear distancing model, the behavior change parameter will be *k* = 3.

### Sequential forecasting with ensemble-based Kalman filter

Sequential forecasting refers to repeated prediction and refinement of state and parameter estimates of the dynamic model based on new information collected about the ground truth. Denote the *n* dimensional vector of unknown state and parameters of a system at time *t* by *x*_*t*_. For instance, in the SIR model for a fixed total population, we can have *x*_*t*_ = (*S*_*t*_/*N*, *I*_*t*_/*N*, *N*, *β*, *δ*, *k*) where *S*_*t*_/*N* and *I*_*t*_/*N* represent the fraction of susceptible and infected individuals at time *t*, respectively. Suppose we make noisy observations about the system’s state at times *t* = 1, 2, …. The observation at time *t* denoted by *y*_*t*_ is an *m* dimensional vector. For instance, weekly noisy measurements may be available for the fraction of infected in the SIRS model, i.e., *y*_*t*_ = *I*_*t*_/*N* + ω_*t*_ where ω_*t*_ is an additive measurement noise. Estimates of the latent states can be updated by using Bayes’ rule, that is, *P*(*x*_*t*_|{y_s_}_*s*=1,…,*t*_) ∝ *P*(*x*_*t*_|{*y*_*s*_}_*s*=1,…,*t*−1_)*P*(*y*_*t*_|*x*_*t*_) where *P*(*x*_*t*_|{*y*_*s*_}_*s*=1,…,*t*_) is the posterior distribution, *P*(*x*_*t*_|{*y*_*s*_}_*s*=1,…,*t*−1_) is the prior distribution, and *P*(*y*_*t*_|*x*_*t*_) is the likelihood of the observation given the prior state. The Bayes’ update rule quickly becomes computationally intractable when the cardinality of the state increases for most prior and noise distributions and nonlinear system dynamics. The computational intractability of the Bayes’ rule has led to development of filtering algorithms that approximate the posterior computation of the Bayes’ rule such as particle filtering^29^, the simple Kalman filter^30^ or the ensemble-based Kalman filter (EnKF)^31,32^.

The ensemble-based Kalman filter (EnKF) is a Monte-Carlo sampling based implementation of two steps: prediction and correction. Specifically, EnKF begins with *M* realizations of the state space *x*_*t*_. We represent realization *i* of the *n* dimensional state at time *t* − 1 by the *n* dimensional vector 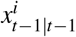, and the ensemble vector array at time *t* − 1 by 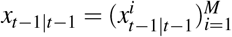.In the prediction step, we integrate each state realization 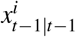 using the model dynamics, e.g., SIR dynamics (1)–(3), to get the prediction 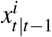. The predicted ensemble mean and covariance are given by

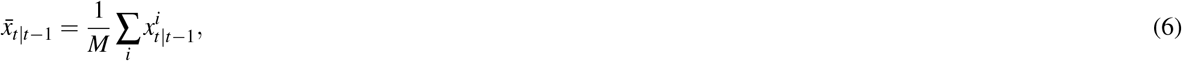

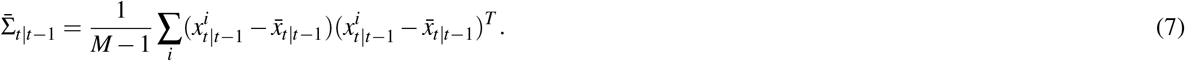

In the correction step, we use 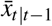 and 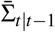 to compute the mean and covariance estimates of the observation *y*_*t*_, denoted by *ŷ_*t*_* and *Ô_*t*_* respectively. For instance, if *x*_*t*_ = (*S*_*t*_/*N*, *I*_*t*_/*N*, *N*, *β*, *δ*, *k*) and *y*_*t*_ = *I*_*t*_/*N* + ω_*t*_ where ω_*t*_ is additive white noise term, the second element of 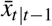 is the observation prediction ŷ_*t*_. We obtain the corrected ensemble by adjusting the ensemble predictions as follows

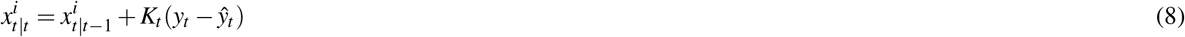

where *K*_*t*_ is an *n* × *m* gain matrix. The gain matrix *K*_*t*_, also known as the Kalman gain, gauges the importance of the measurement relative to the prediction. Formally, it minimizes the mean square error of the estimate and can be solved by solving a set of linear equations—see SI Section B. The steps described above are repeated after each observation.

In the initialization of EnKF, we select the ensemble size *M*, and sample *M* values from a prior distribution of the state space. We denote the *i*th realization of the state space by 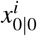. Further, in our implementation of the EnKF, we add zero-mean noise to the predictions of the observation. That is, instead of a single prediction of the observation ŷ_*t*_ for the entire ensemble, we have a prediction for each ensemble member 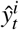 for all *i* = 1, …, *M*—see SI Section B for details. Adding an artificial noise to our predictions prevents ensemble values from getting too close to each other after repeated correction steps^33^.

### Numerical implementation of the ensemble-based Kalman filter

We assume the initial fraction of susceptible and infected (*S*_0_/*N* and *I*_0_/*N*), and the behavior parameter *k* are unknown. In this paper, our focus is on comparing the accuracy of sequential forecasting with and without behavior change. Hence, we assume *N*, *β* and *δ* are known. Then the state space of EnKF is given by the 3×1 vector *x*_*t*_ = (*S*_*t*_/*N*, *I*_*t*_/*N*, *k*). We will have *M* = 20 samples in an ensemble, unless explicitly stated otherwise. We denote the 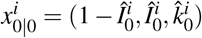. Initial state realizations of the fraction of infected come from a Gaussian prior distribution with mean*I*_0_/*N* and variance 0.05, that is, let 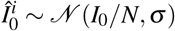. If the prior value realized is negative, we discard it, and redraw from the Gaussian prior distribution until all realizations of 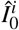 are non-negative. We will specify the prior distribution on 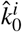 based on the specific scenarios considered in the Results section. We assume there are weekly noisy measurements of the fraction of infected, *y*_*t*_ = *I*_*t*_/*N* + *w*_*t*_, where *t* = 1, 2, … weeks and *w*_*t*_ is a Gaussian random variable with zero mean and variance 10^−3^ that indicates measurement noise level. Given the prior ensemble vector *x*_*t*−1|*t*−1_, we construct the predicted ensemble vector *x*_*t*|*t*−1_ by integrating the SIR dynamics in (1)–(3) for 1 week. We obtain the mean measurement prediction by taking the mean of the second element of the ensemble members, that is, 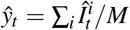. We add prediction noise to ŷ_*t*_ during the update of each ensemble member in (8). We select prediction noise to be zero-mean Gaussian with a variance that we specify in Results section based on the scenario.

## A The final outbreak size of the long-term awareness model

In the long-term awareness model we can simplify the distancing term as 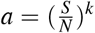. The ratio of change in susceptible individuals to the change in recovered individuals is given by

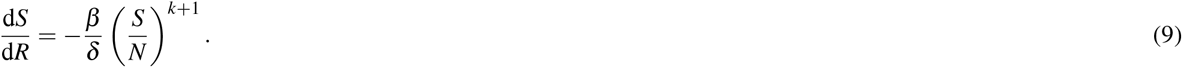

For no social distancing (*k* = 0), we recover the well-known infinite time relationship for susceptible individuals^22^:

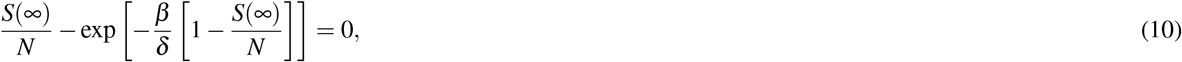

where we simplified the result of integration by assuming *R*(0) = 0 and *S*(0) ≈ *N*, and noting that *R*(∞) = *N* − *S*(∞). There does not exist a closed form solution for *S*(∞)^34^. In Figure 1, we plot the above function on the left hand side and show critical *S*(∞) that equates it to zero (see case *k* = 0).

For *k* > 0 where individuals modify their behavior, we obtain the following equation by integrating (9) from time zero to infinity,

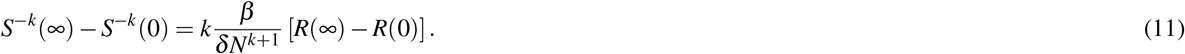

Assuming *R*(0) = 0 and *S*(0) ≈ *N*, and noting that *R*(∞) = *N* − *S*(∞) since *I*(∞) = 0, we have

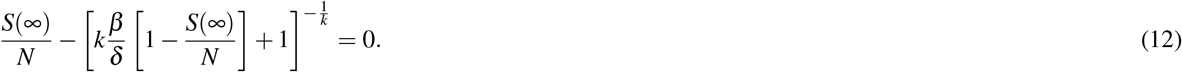

Solving the above equation when *k* = 1 gives the closed form solution that

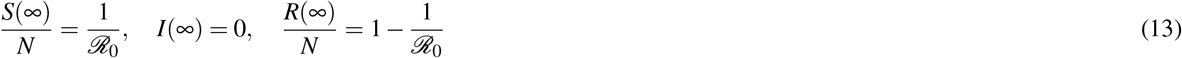

where 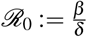.

## B Observation prediction and Kalman gain

Let the observation at time *t y*_*t*_, an *m* × 1 vector be given by

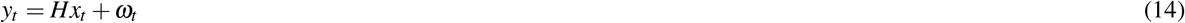

where *H* is the *m* × *n* observation matrix and ω_*t*_ is the measurement noise vector of size *m* × 1. The observation prediction of the *i*th member of the ensemble is given by

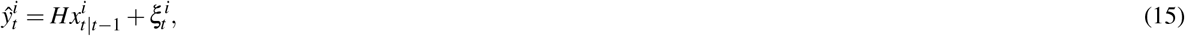

where 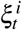 is the zero-mean prediction noise added to each ensemble prediction. We can compute the predicted mean and covariance of the observation as follows

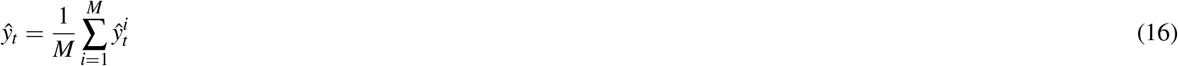

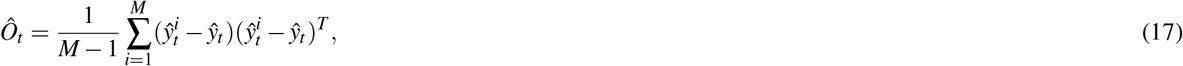
where *Ô*_*t*_ is an *m* × *m* matrix. Next, we compute the predicted cross-covariance between the state and observation, an *n* × *m* matrix, which is also known as the innovation covariance,

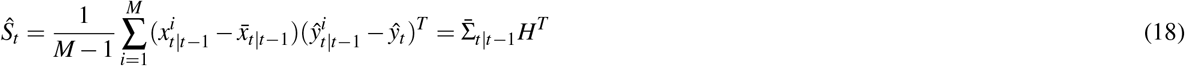

where 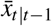 is defined in eq. (6). The Kalman gain *K*_*t*_, an *n* × *m* matrix, minimizes the variance by solving the following set of equations,

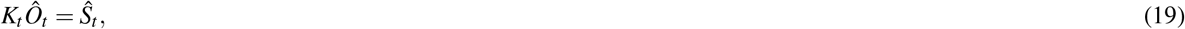

which means we can compute *K*_*t*_ = *Ŝ*_*t*_ *Ô*_*t*_^−1^.

## C Effect of external and algorithm parameters on sequential forecasts

The external parameters are the prior distributions of the state space, e.g., *k* and *I*_0_ prior estimates, the distribution of the measurement noise, and other model parameters. In the main text, we assumed the initial estimate of the behavior parameter is Gaussian with mean *k̅* = 1 and variance 0.5. Here, we vary the mean of the prior distribution *k̅* and analyze its effect on forecast error in Fig. 5. Fig. 5 shows that when the prior mean estimate is closer to the ground truth, our initial forecast is better. Specifically, the forecast error reduces more than half when the prior mean estimate changes from *k̅* = 1 to *k̅* = 2. Regardless of the prior mean, forecast errors converge to similar forecast error values by the 20th week. This shows that while the prior mean affects the early forecast errors, it does not affect the learning rate. The learning rate depends on our confidence in the prior estimate and the measurement noise, that is, the prior variance and measurement variance. Along these lines in Fig. 6, we show that having a high variance in the prior is better if our initial mean estimate of the state is not close to the true value where we measure expected forecast error for different prior mean (*k̅*) and variance values. The reasoning is that when the mean estimate is correct, it is better to have high confidence in our estimate. However, when the mean estimate is wrong, the confidence in the estimate should be low so that our initial ensemble sample have realizations close to the ground truth.

**Figure 5.**
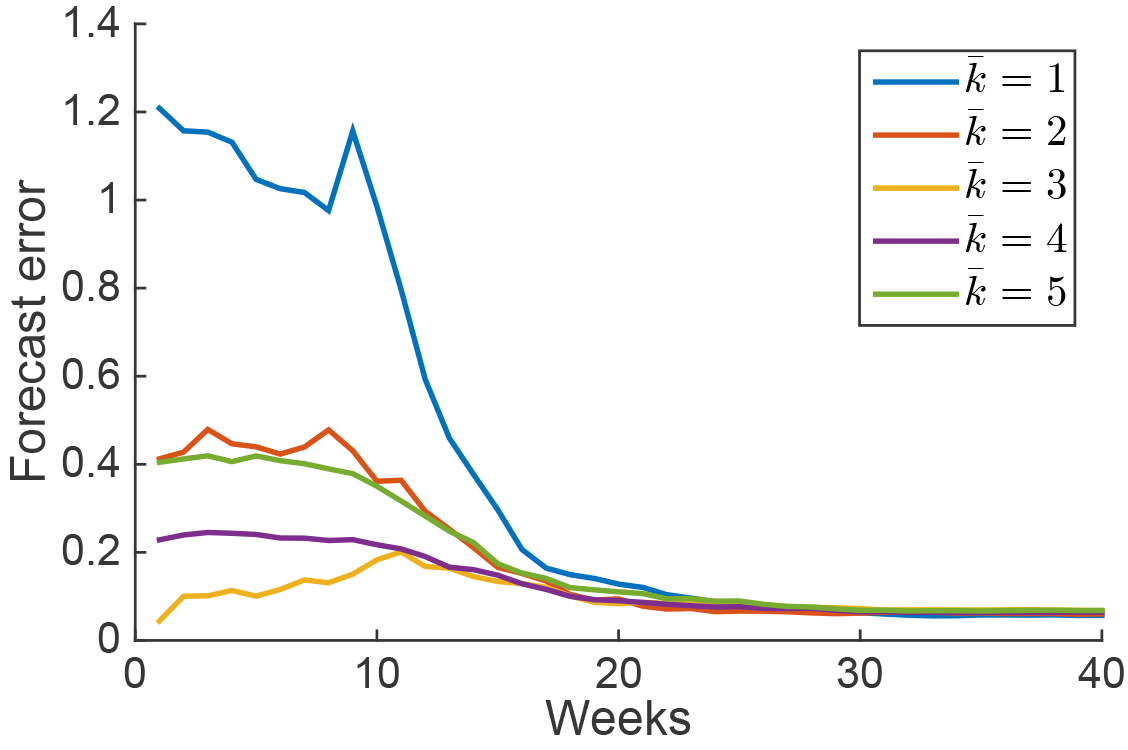
Effect of prior mean estimate of the behavior parameter (*k̅*) on forecasts. We consider the setup of the second sequential forecasting scenario with EnKF shown in Fig. 3(b) with long-term awareness model. The ground truth behavior parameter is *k* = 3. Prior distribution of *k* is Gaussian with mean *k̅* and variance equal to 0.5, that is, 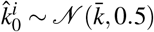. We vary *k̅* ∊ {1,2,3,4,5}. For the estimated ensemble at time *t*, 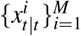, we construct trajectories 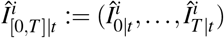 for the number of infected by integrating 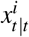 forward and backward. The mean trajectory of the ensemble is then the 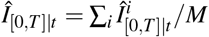. The forecast error of the ensemble is the norm of the difference between the ensemble trajectory (*Î*_[0,*T*]|*t*_) and the ground truth trajectory (*I*_[0,*T*]|*t*_), i.e., ‖*Î*_[0,*T*]|*t*_) − *I*_[0*,T*]*|t*_‖. Initial forecast errors show the value of the prior mean estimate. When the prior mean is correct, i.e., when *k* = *k*, the initial forecast error is the smallest. After the 20th week, forecast errors are small for all prior means.

**Figure 6.**
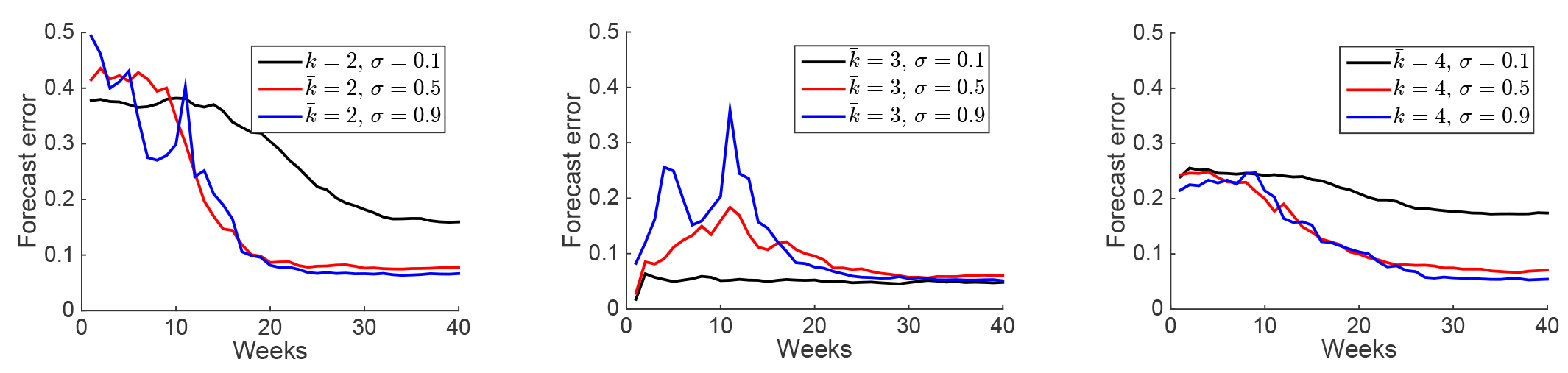
Forecast error with respect to prior distribution on behavior parameter. The ground truth behavior parameter is *k* = 3. We consider the numerical setup in Fig. 3 (b) with long-term awareness model. We sample *M* = 20 initial ensemble values of the behavior parameter 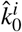 according to a Gaussian distribution with mean *k̅* and variance σ. Small variance implies high certainty on our mean estimate. When the prior mean estimate is not correct *k̅* ≠ *k*, small variance worsens performance—see left (*k̅* = 2) and right (*k̅* = 4) with σ = 0.1. When the prior mean estimate is correct *k̅* = *k*, small variance improves performance—see center (*k̅* = 3).

When the observation noise level decreases, the EnKF performance improves as can be expected—see Fig. 7. The EnKF parameters include the sample size *M*, the sampling distribution of the initial ensemble, and the distribution of the prediction noise. An important question is how to select the EnKF parameters given the external parameters so that our forecasts get close to the ground truth faster. When the sampling size *M* is small(*M* ≈ 10), the sequential forecasting with EnKF performs poorly—see Fig. 8.

**Figure 7.**
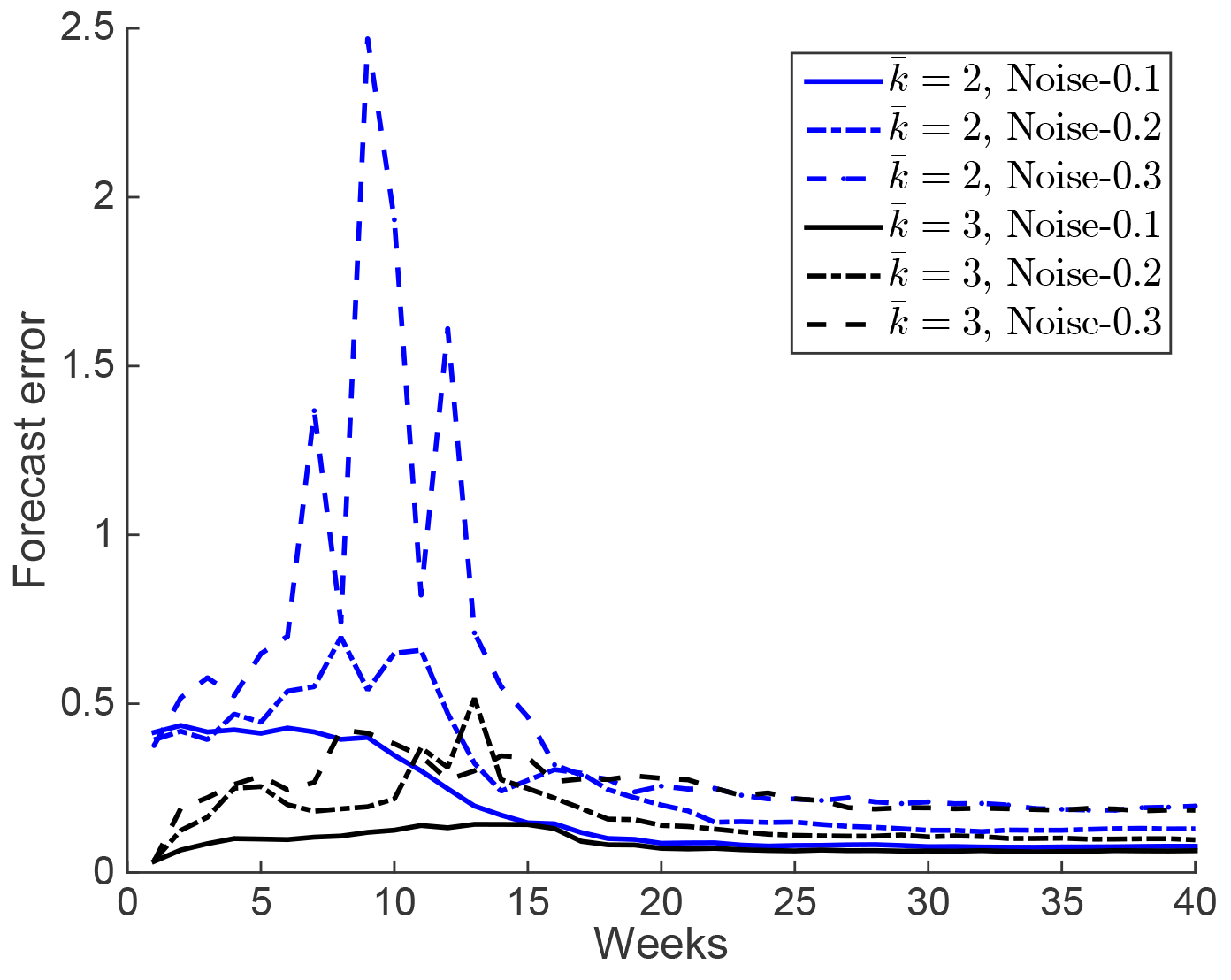
Effect of measurement noise on forecast errors. We consider the numerical setup in Fig. 3(d). Measurement noise is zero mean with variance that is selected from the set {0.1,0.2,0.3}. We measure the forecast error as defined in Fig. S1. Lower measurement noise improves EnKF.

**Figure 8.**
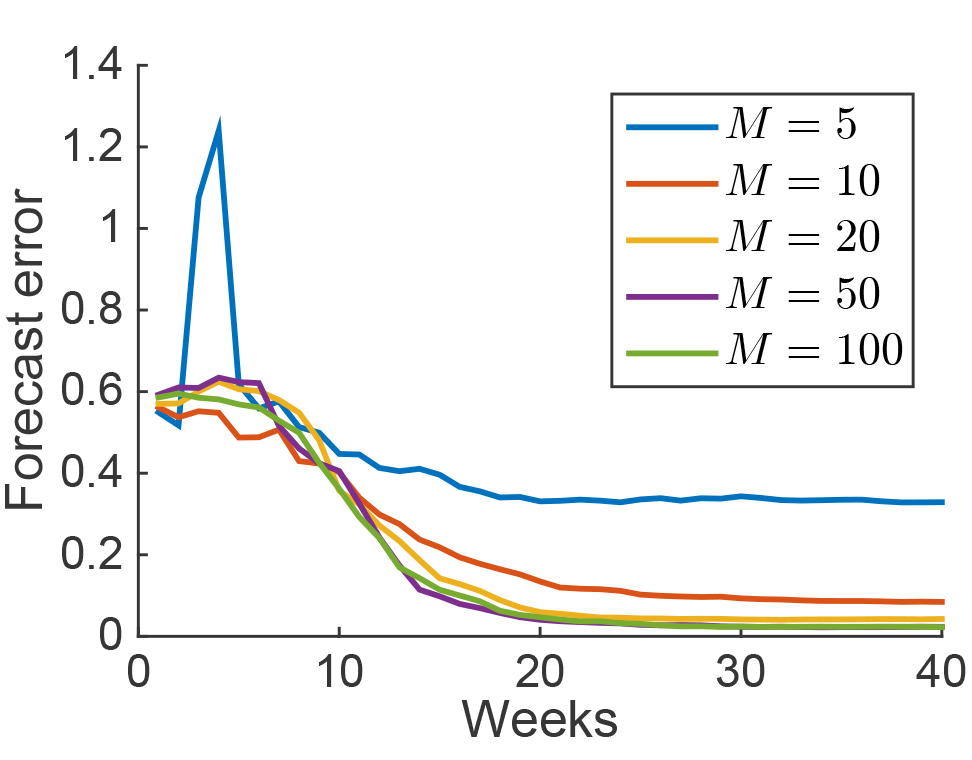
Effect of ensemble size on forecast errors. We consider the numerical setup in Fig. 3(d). We measure the forecast error as defined in Fig. S1. We observe diminishing improvement in forecast errors as we increase the ensemble size. As ensemble size (*M*) increases from 5 to 20, the accuracy of forecasts improve significantly. As ensemble size increases from 20 to 50 and 100, the improvement is less significant.

Finally, recall that we add artificial prediction noise to each sample to avoid the ensemble values getting too close to each other early. Our experimentation with the prediction noise distributions showed that the prediction noise values that achieve a good tracking of the ground truth in weekly estimates, i.e., where 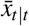 is close to *x*_*t*_, depend on the scenario. For the first scenario (prediction without behavior), we find that the prediction noise variance has to be smaller than 10^−8^ but positive for good tracking—see Fig. 9. For the second scenario (prediction with unknown behavior parameter), the posterior estimates are good when the prediction noise variance is between 10^−4^ and 10^−3^−see Fig. 10. In Fig. 3, we select the variance value that minimizes the forecast error for each scenario—see Fig. 11. These results imply that the performance of the EnKF method is sensitive to the selection of prediction noise. Further, the performance depends on the structural accuracy of the model. When the model is inaccurate, smaller prediction noise values give better tracking performance but still fail to predict the future (see Fig. 3(a) and (c)). When the model is accurate, we need to select the prediction noise smaller than the observation noise but not too small.

**Figure 9.**
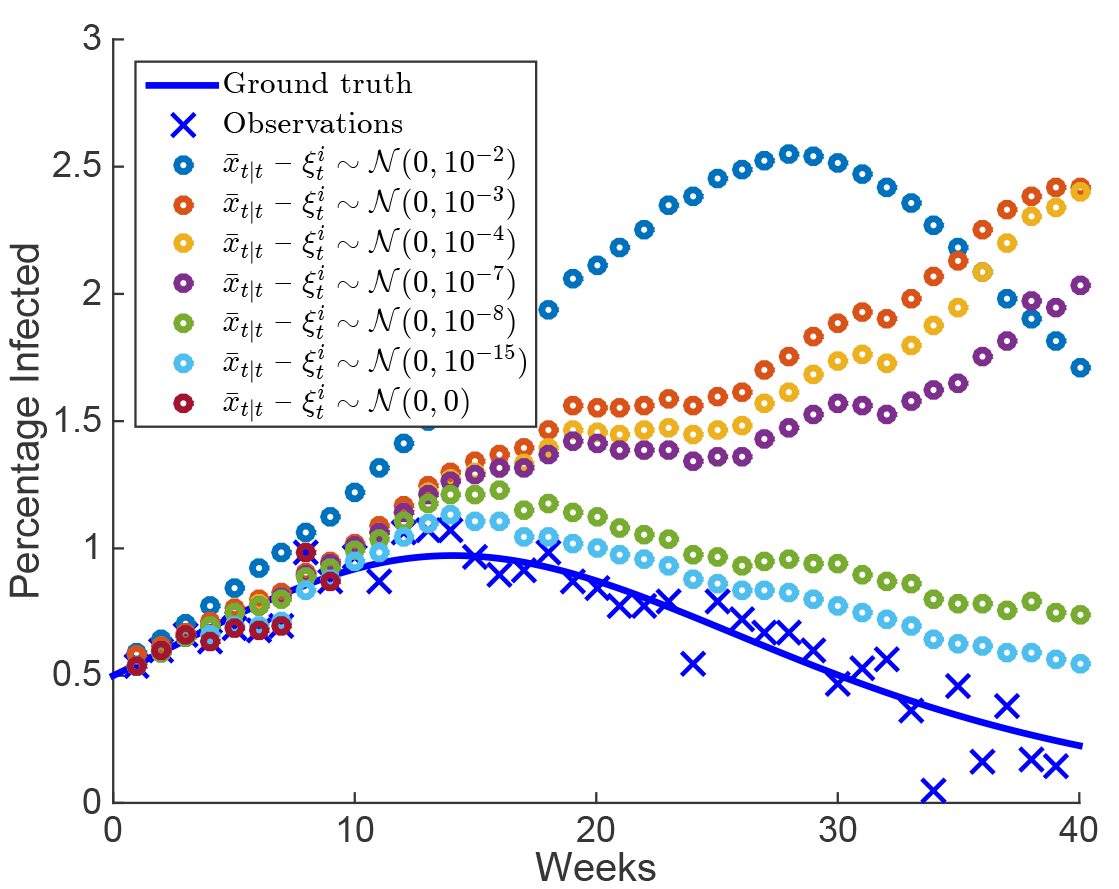
The effect of prediction noise variance on sequential estimates of a synthetic outbreak with long term awareness with an inaccurate model. We consider the numerical setup in Fig. 3 (c). We set the variance of the prediction noise 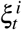 in (15) equal to {10^−2^,10^−3^,10^−4^,10^−7^,10^−8^,10^−15^,0}. Each point is the ensemble’s weekly mean estimate of the percentage of infected, that is, the second element of 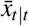 in eq. (8). Weekly estimates track the ground truth better when the variance is smaller than 10^−8^. When the variance is zero, tracking fails after Week 10 because the ensemble values get very close to each other causing a division by zero in computing the Kalman gain (19).

**Figure 10.**
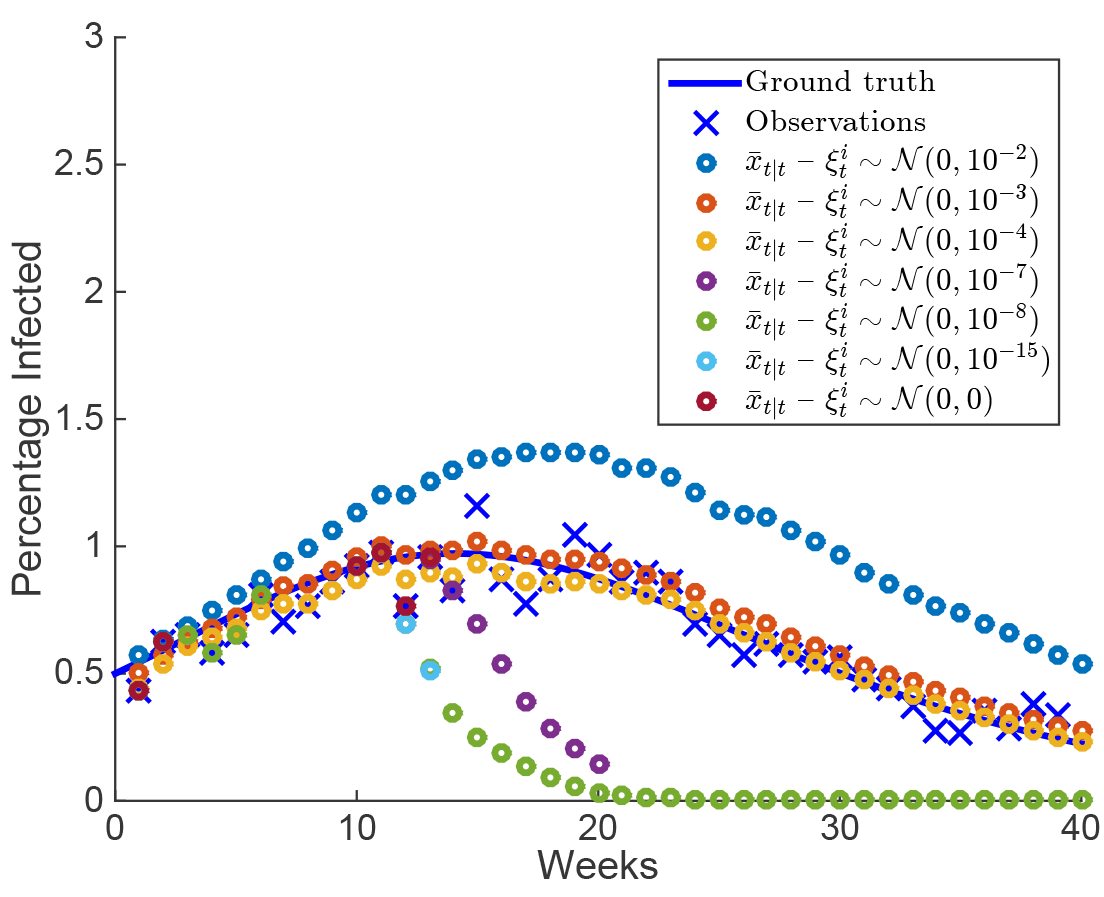
The effect of prediction noise variance on sequential estimates of a synthetic outbreak with long term awareness with a model that accounts for behavior change. We consider the numerical setup in Fig. 3 (d). We set the variance of the prediction noise 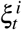 in (15) equal to {10^−2^,10^−3^,10^−4^,10^−7^,10^−8^,10^−15^,0}. Each point is the ensemble’s weekly mean estimate of the percentage of infected, that is, the second element of 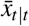 in eq. (8). Weekly estimates track the ground truth well when the variance is 10^−3^ or 10^−4^,. When the variance is larger than 10^−3^, the estimates are erroneous. When variance is 10^−7^, or smaller, the EnKF computations fail after certain weeks or become meaningless. That is for variance smaller than 10^−7^, we have premature convergence of the ensemble values causing a division by zero in computing the Kalman gain (19).

**Figure 11.**
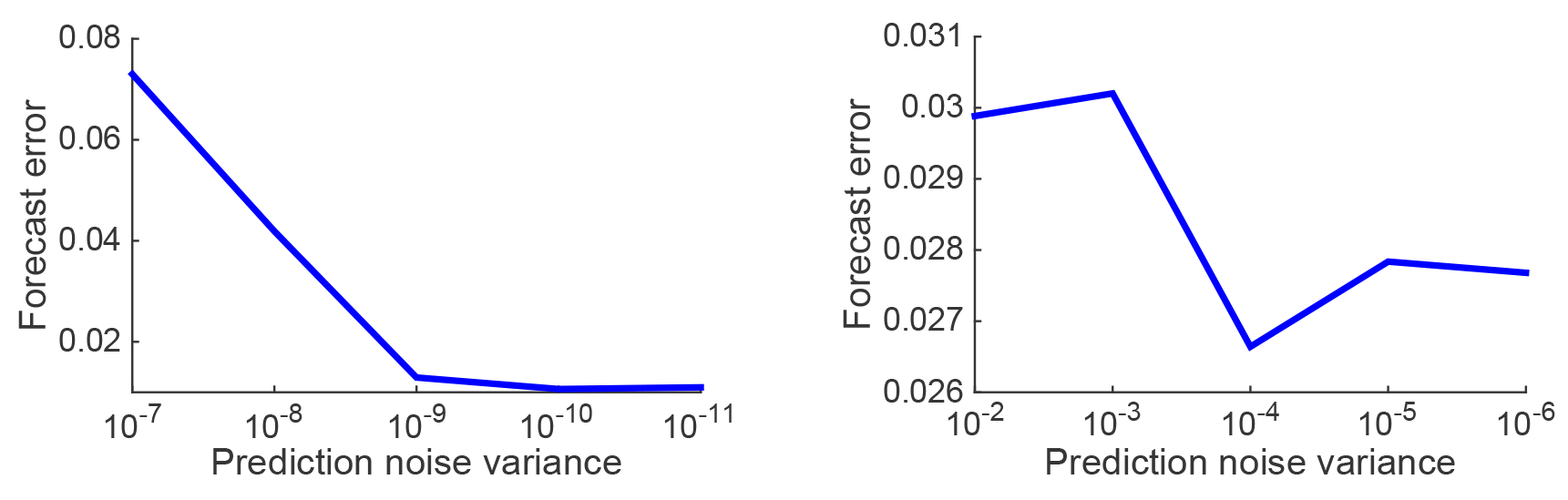
Forecast error with respect to prediction noise variance. Prediction noise in (15) is distributed according to zero mean Gaussian with a given variance. For each variance value, we run the EnKF algorithm for multiple realizations of the epidemic. For each run, we compute the forecast error of the weekly estimates. The forecast error value shown here for each variance value is an average of the forecast errors of the multiple runs. (Left) EnKF predictions that do not account for behavior change—first scenario. (Right) EnKF predictions that account for behavior change—second scenario. The minimum forecast error is 10^−11^, and 10^−4^, respectively for Left and Right. For the first scenario (left), the forecast error continues to decrease as we further decrease the variance. In Fig. 3 (a) and (c), we choose the variance to be 10^−10^, because the decrease in forecast error is negligible for values smaller than 10^−10^,.

